# Unravelling Biosynthesis and Biodegradation Potentials of Microbial Dark Matters in Hypersaline Lakes

**DOI:** 10.1101/2023.06.28.546814

**Authors:** Zhiguang Qiu, Yuanyuan Zhu, Qing Zhang, Xuejiao Qiao, Rong Mu, Zheng Xu, Yan Yan, Fan Wang, Tong Zhang, Wei-Qin Zhuang, Ke Yu

## Abstract

Biosynthesis and biodegradation of microorganisms critically underpin the development of biotechnology, new drugs and therapies, and environmental remediation. However, the vast majority of uncultured microbial species along with their metabolic capacities in extreme environments remain obscured. To unravel the metabolic potentials of these microbial dark matters (MDMs), we investigated four deep-inland hypersaline lakes with largely diversified environmental parameters in Xinjiang Uygur Zizhiqu, China. Metagenomic binning obtained 3,030 metagenome-assembled genomes (MAGs) spanning 82 phyla, of which 2,363 MAGs could not be assigned to a known genus. These unknown MAGs were abundantly observed with distinct taxa among lakes, possibly linked to the diversification of physiochemical conditions. Analysis of biosynthetic potentials identified 9,635 biosynthesis gene clusters (BGCs), of which 9,403 BGCs were considered novel. We found that some MAGs from putatively novel phyla consistently comprised enriched BGCs, which may have substantial potentials in biotechnological applications. In addition, biodegradation potentials such as dehalogenation, anaerobic ammonium oxidation (Anammox), polycyclic aromatic hydrocarbon (PAH), and plastic degradation were found in new microbial clades from hypersaline lakes. These findings substantially expanded the genetic repository of biosynthesis and biodegradation potentials, which can further assist the development of new and innovative applications in biotechnology.

## 1. Introduction

Microorganisms are vital in global ecosystems that underpin life on Earth, as they play essential roles in contributing to biogeochemical cycling, supporting food webs, and maintaining the fitness of plants and animals [1–4]. Particularly, the microbial metabolic capabilities such as decomposing pollutants and producing secondary metabolites like natural products have fostered the development of biosynthesis and biodegradation, thereby propelled biotechnological processes and addressed environmental challenges [5–8]. Nonetheless, most microbial species are unculturable, leading to the inefficient exploration of metabolic capacities across diverse environments.

To address this issue, a series of state-of-the-art sequencing technologies along with bioinformatic tools, such as metagenomic sequencing and binning, have revolutionised our ability to investigate and decode the metabolic pathways of uncultured microorganisms. Employing such approaches, we can unveil previously uncharacterised microorganisms within an underexplored environment, considerably expanding our recognition of the specific microbial roles and functions in biosynthesis and biodegradation. This knowledge could be instrumental in harnessing more efficient and sustainable methods for producing natural products or devising more effective pathways for contaminant degradation [9]. However, current progress in applying microbial resources for biosynthesis is still hindered by the efficiency of the enzymes for large-scale production, the sustainability of methodologies, and the financial demands of culturing [10, 11]. On the other hand, environmental variability, the complexity of microbial communities, and the high pre-treatment costs are the main issues that obstructed biodegradation efficiency [12, 13]. Furthermore, our understanding of the underlying microbial mechanisms of biosynthesis and biodegradation, including novel enzymes and metabolic pathways involved in these processes, remain considerably limited. Therefore, to enhance our understanding of these mechanisms and facilitate more efficient and effective biosynthesis and biodegradation processes, it is imperative to investigate uncultured microbial species and their metabolic capacities in extreme environments. Such exploration can deepen the knowledge of novel enzymes and metabolic pathways, leading to more sustainable and environmentally friendly solutions across various applications.

Hypersaline lakes are typical examples of extreme environments, usually characterised by high salinity (≥ 35 g L^−1^), elevated aridity and evaporation, poor nutrients such as organic carbon, and some even exhibiting high alkalinity [14–16]. Within these harsh conditions, unique extremophilic microbial communities have thrived, showcasing an exceptional ability to acclimate to environments inhospitable for other living organisms [17, 18]. The distinctive environmental parameters of deep inland hypersaline lakes vary significantly from one lake to another, shaped by climate, geology, and human activities. Consequently, many microbial communities in these hypersaline habitats remain unexplored, presenting potential resources for biotechnological applications [19–22]. Genome-resolved analyses indicates that these deep inland hypersaline lakes many uncultured microorganisms, namely microbial dark matters (MDMs), such as DPANN archaea and CPR bacteria [15, 23, 24]. These findings substantially broadened our recognition of MDMs from an evolutionary perspective. However, these pioneering studies merely scratched the surface of hypersaline lake microbiomes, restricted by the scope of study areas and limitations in bioinformatic analysis techniques. Therefore, to comprehensively uncover the microbial niches within hypersaline lake microbiomes and further reveal metabolic capacities in these extreme ecosystems, more in-depth and expansive investigations on hypersaline lake microbial genomes should be envisaged.

In this study, we examined four saline lakes in Xinjiang Uygur Zizhiqu, China, each with distinct salinity (1.5–237 g L^−1^) and altitude (−153 to 1,585 m). A previous study observed that salinity significantly influences the microbial assemblages of saline lake microbiota [25]. However, the roles lakewater and sediment microbial communities devote to biosynthesis and biodegradation resolved from metagenomes remain uncertain. Given such a diverse range of geographic and environmental parameters, we assembled over 3,000 metagenome-assembled genomes (MAGs) from these four lakes, aiming to unearth new uncultured microbial lineages and decode the biosynthetic and biodegradative potentials of hypersaline lake prokaryotes.

## 2. Methods

### 2.1. Sampling and measurement of physiochemical parameters

Hypersaline lake samples were collected in July 2018 from Aiding Lake (ADH), Barkol Lake (BLK), Dabancheng Lake (DBC), and Qijiaojing Lake (QJJ) (Fig. S1), Xinjiang Uygur Zizhiqu, China. Detailed sampling and geographic information were recorded in Table S1. Briefly, four water samples (*n* = 4) and four sediment samples (*n* = 4) were collected from each lake, at least 50 m apart between two sampling sites. For each water sample, 5 L of lake water from the upper 50 cm surface was randomly collected into a 5 L sterile container. Lake water was firstly filtered through a filter paper (Whatman, GE Healthcare, NY, USA) to remove large particles before filtering through 0.22 μm PES (Polyethersulfone) membranes (Millipore, Billerica, MA, USA) to enrich lakewater microorganisms. For each sediment sample, about 50 grams of sediments were randomly collected at 0–10 cm depth at the bottom of the lake into sterile 50 mL falcon tubes. Membranes and sediment samples were immediately placed on dry ice before being brought to the laboratory and transferred to a –80 °C freezer until further DNA extraction was performed.

The salinity and pH of lake water were measured *in situ* using a Hydrolab DS5 multiparameter water quality meter (Hach Company, Loveland, CO, USA). Total organic carbon (TOC) was analysed with a TOC/TN-VCPH analyser (Shimadzu, Tokyo, Japan). The concentrations of lithium (Li^+^), sodium (Na^+^), ammonium (NH_4_^+^), potassium (K^+^), magnesium (Mg^2+^), calcium (Ca^2+^), chloridion (Cl^−^), and sulphate (SO^42−^) were determined using an inductively coupled plasma mass spectrometry (Agilent Technologies Inc., Bellevue, WA, USA).

### 2.2. DNA extraction and metagenomic sequencing

Total DNA was extracted from frozen filtered membrane or sediment samples using the DNeasy PowerSoil Pro Kit (Qiagen, Hilden, Germany), following the manufacturer’s instructions. Extracted DNA was quality checked by NanoDrop 2000 (Thermo Fisher Scientific, Waltham, Massachusetts, US) and quantity checked by Qubit Fluorometer (Thermo Fisher Scientific, USA). Next-generation metagenomic sequencing was performed at Novogene (Tianjin, China) on an Illumina NovaSeq 6000 platform using the PE150 strategy.

### 2.3. Assembly, binning, and phylogenetic analyses

Raw paired-end reads were initially filtered using fastp with default parameters [26]. Clean reads were individually and co-assembled using SPAdes (v3.15.2) [27] under “--meta” mode into contigs specifying k-mer sizes of 21, 33, 55, 77, and finally reserved contigs > 1,000 bps. Metagenomic binning was applied to both single sample assembly and co-assembly using BASALT (v1.0) under default mode [28]. The completeness and contamination of bins were assessed using CheckM v1.0.2 with “lineage_wf” workflow and default parameters [29], with only medium and high-quality bins that meet the MIMAG standard (completeness ≥ 50, contamination ≤ 10) [30] kept as MAGs for further analysis.

Taxonomic classification was conducted using GTDB-Tk (v1.5.0, database release r202) [31]. MAGs not classified to any reference genomes in GTDB were defined as unknown at certain taxonomic levels (i.e., species, genus, family, etc.). Phylogenetic analysis of bacteria and archaea MAGs was performed based on a multiple sequence alignment of 120 bacterial- and 122 archaeal-specific single-copy marker proteins, respectively. A concatenated alignment of these marker proteins was created using HMMER v3.1.b2 with default parameters [32]. Bacterial and archaeal phylogenetic trees were inferred using IQ-TREE under the best-fitted models with 1,000 bootstrap replications [33] before being visualised and annotated in iTOL v6 [34].

### 2.4. Identification of non-coding RNA genes and functional annotation

Non-coding RNA genes, including transfer RNA genes and ribosomal RNA genes, were identified to evaluate the integrity of MAGs. Briefly, tRNA genes were predicted using tRNAscan-SE (v2.0.9) [35], while rRNA genes, including 5S rRNA, 16S rRNA, and 23S rRNA, were predicted using Barrnap (v0.9, https://github.com/tseemann/barrnap). For genetic analysis, Opening Reading Frames (ORFs) of assembled contigs were predicted using Prodigal (v2.6.3) with default parameters [36]. Predicted ORFs were dereplicated with CD-HIT (v4.6) at 90% identity [37]. Dereplicated ORFs were annotated against the Kyoto Encyclopedia of Genes and Genomes (KEGG) database (version 58) and National Center for Biotechnology Information – Non-redundant (NCBI-NR) database (version 20220806), respectively, with Diamond (v2.0.11.149) [38] using “more-sensitive” mode and e-value at 1×10^-10^. In addition, dereplicated ORFs were further annotated against Pfam (v34.0) and TIGRFAM (v15.0) databases using InterProScan (v5.54-87.0) [39]. To estimate the relative abundance of each gene, dereplicated FASTA sequences of ORFs were mapped against raw sequence reads by Bowtie2 (v2.4.2) [40]. Then, the depth of aligned reads was calculated using SAMtools (v1.7) [41] and summarised with a Perl script ‘calc.coverage.in.bam.depth.pl’ from a previous study [42]. Finally, coverage-normalised KOs were clustered into modules and pathways using a Python script “pathway_pipeline.py” in PICRUSt2 [43]. For genome-centric analysis, each MAG was predicted using Prodigal with default parameters and annotated against the KEGG database using the same parameter described above.

### 2.5. Annotation of Biosynthetic Gene Clusters (BGCs)

Genome sequences were used as input using antiSMASH (v.6.1.1) [44] to identify BGCs from saline lake MAGs. The gbk-formatted output of each MAG was further processed using BiG-SCAPE (v1.1.2) [45] to cluster saline lake BGCs with reference BGCs from the MIBiG database [46]. Gene Cluster Families (GCFs) were clustered at 0.2, 0.3, 0.4, 0.5, 0.6, 0.7, and 0.8 distance thresholds, respectively. To determine the novelty of BGCs, we followed the criteria described by Navarro-Muñozet et al. [45] using 0.5 distance for clustering. Any BGC not related to MIBiG BGCs were described as unknown BGC.

### 2.6. Detection of Biodegradation potentials

To identify MAGs with potential biodegradation functions, including dechlorinating, nitrogen removal, and degradation of polycyclic aromatic hydrocarbons (PAHs), dereplicated MAG ORFs were further annotated against two homebrew databases, including EDB-DHG and EDB-Ncyc for dechlorinating and nitrogen removal. Concurrently, Darhd Database and PlasticDB were employed to assess PAH degradation [47] and plastic degradation [48], respectively. The annotation process utilised Diamond with the same parameter described above, reserving matched sequences at identity ≥ 50%. As the Darhd database only contained nucleotide sequences, we pre-treated all sequences by translating them into amino acids with Prodigal to maintain consistency with other databases. Annotated sequences were checked by comparing results with KEGG, NCBI-NR, and InterProScan to avoid false annotation. EDB-DHG and EDB-Ncyc Databases are available for download in FigShare (https://figshare.com/ndownloader/articles/23504874/versions/1).

### 2.7. Statistical analysis and data visualisation

Data organisation and formatting were conducted using the R packages “tibble”, “dplyr”, “tidyr”, and “stringr”. Statistical analyses, including Kruskal-Wallis tests, Dunn tests, and RDA analysis, were conducted using the R packages “vegan”, “stats”, and “FSA”. Data visualisation, including dot plots, bar graphs (including pie charts), and boxplots, were performed using the R package “ggplot2” with functions “geom_point”, “geom_bar”, and “geom_boxplot”, respectively. nMDS plots and RDA plots were performed using the R packages “ggplot2” with functions “geom_point”, “geom_hline”, “geom_vline”, and “geom_segment”, respectively. Heatmap was generated using “pheatmap” with z-score normalised using “scale = row”, and multiple panels were generated using “cowplot”.

## 3. Results

### 3.1. A large proportion of metagenome-assembled genomes from hypersaline lakes were considered novel

Metagenomic sequencing generated ∼1.2 Terabytes of DNA sequence data from 30 samples from four hypersaline lakes. Two samples (one from DBC water and the other from QJJ sediment) were excluded from this study due to the unsuccessful preparation of the sequencing library. The metagenomic assembly generated 26 Gigabytes of contig data with sequence length > 1,000 bp. After metagenomic binning, we recovered 3,030 non-redundant bins that meet the MIMAG standard (completeness ≥ 50%, purity ≥ 90%, mean completeness = 79.8 ± 14.8%, mean purity = 97.8 ± 2.0%) [30]. These bins were considered as MAGs for the following analyses (Fig. 1a). There were 76.9%, 51.5%, and 44.6% of the MAGs contained 5S, 16S, and 23S rRNA genes, respectively, while 23.7% possessed all three types of rRNA genes (Fig. 1b). In terms of tRNA genes, 75% of the MAGs contained more than 13 types of tRNA genes (Fig. 1c). There were 97.1%, 78.0% and 56.7% of MAGs affiliated with unclassified or close to a tentatively assigned species, genera, and families, respectively, suggesting that a high proportion of the genomes discovered in the hypersaline lakes were unknown or candidates for future identification (Fig. 1d). Details of the sequencing data size, contig summary, number of genomes, and specifics of the MAGs were supplied in Table S1.

**Fig. 1.**
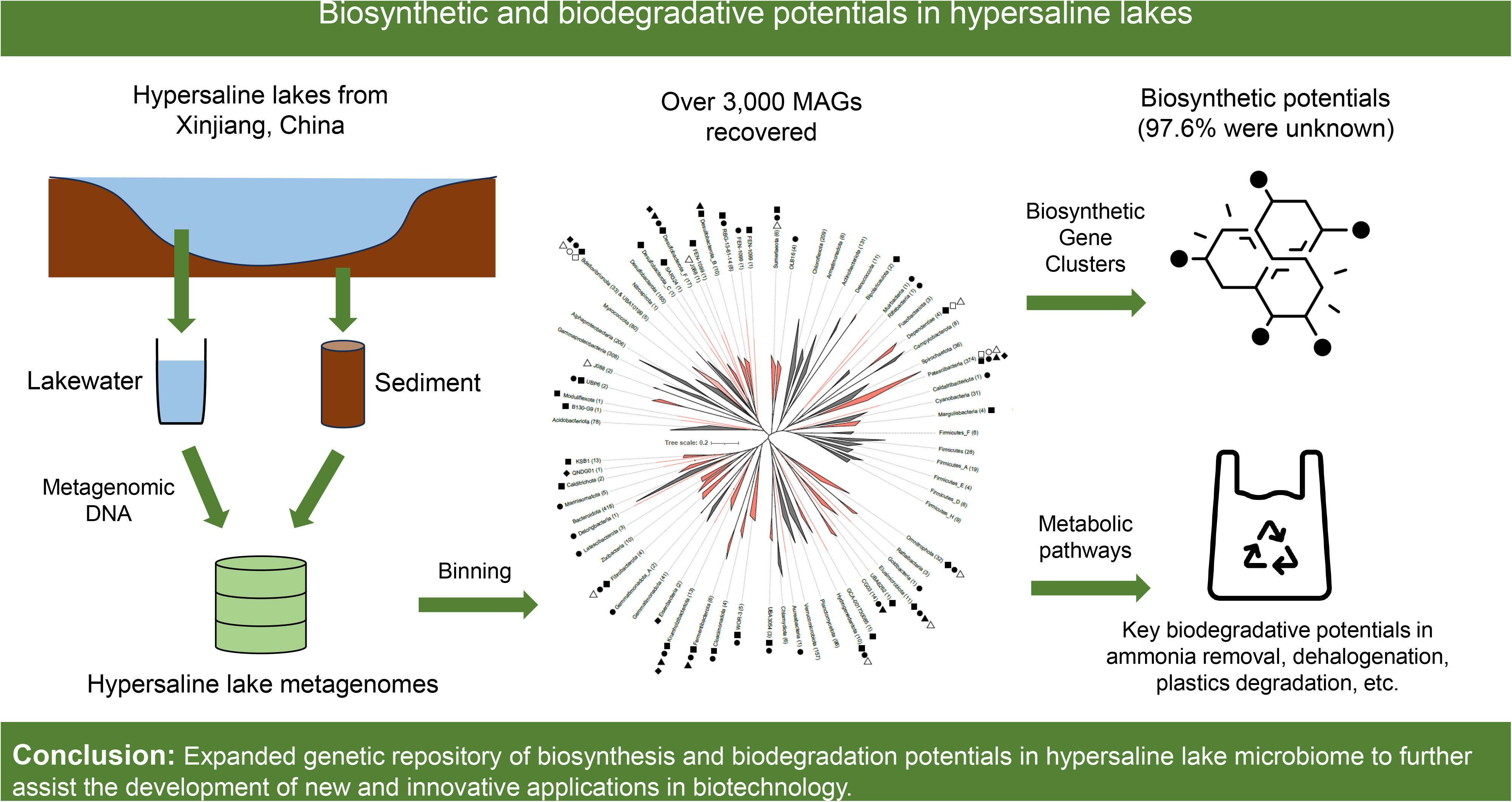
Summary of MAGs obtained from hypersaline lakes. **a**, A breakdown of completeness and purity of recovered MAGs. Red: medium-quality MAGs; Green: high-quality MAGs; Blue: Near-complete (NC) MAGs. **b**, The proportion of 5S, 16S, 23S, and all three types of rRNA genes predicted in MAGs. **c**, Summary of genes encoding standard tRNAs predicted in MAGs. **d**, Proportion of MAGs affiliated with a classified, candidate, or unknown taxon at different levels.

### 3.2. Distinct genetic composition and functional potential among lakes

Among functional genes from all assembled contigs, distinct genetic compositions were identified among the lakes and sample types. Overall, the lakewater genetic richness in the Chao1 index of ADH and DBC was significantly higher than that of QJJ, while the genetic richness and evenness in the Shannon index of DBC was significantly higher than that of QJJ (*P* < 0.05). However, no significant difference was found between lakes in sedimentary samples (Fig. 2a). Genetic compositions of lake lakewater and sedimentary communities were significantly different (*P* < 0.05, Fig. S2), which were mainly driven by salinity including Li^+^, Na^+^, SO_4_^2−^, Ca^2+^, Mg^2+^, and K^+^ in lakewater communities, and TOC, SO_4_^2−^, and Na^+^ in sedimentary communities (Fig. 2b). Functional capacity profiles suggested that microbial genetic potentials varied distinctly among sample types and lakes, particularly in aspects such as nitrogen and sulphur metabolisms, carbon fixation, methanogenesis, and multidrug resistance, among others (Fig. 2c). Specific analysis within pathways showed that in carbon fixation, Dicarboxylate−hydroxybutyrate cycle was mainly enriched in QJJ planktons, while rTCA cycle, Hydroxypropionate−hydroxybutylate cycle, and 3−Hydroxypropionate bi-cycle were enriched in QJJ sediments. On the other hand, BLK sediment communities mainly performed the Reductive pentose phosphate cycle (CBB cycle) and Wood−Ljungdahl pathway for carbon fixation. For methane metabolism, BLK was found to be enriched in Methanogenesis in sediments, while DBC and QJJ were enriched in methane oxidation in sediments. F420 biosynthesis was found to be abundant in QJJ plankton, possibly due to the high relative abundance of archaea communities. Nitrogen cycle-related modules, including nitrogen fixation, nitrification, and denitrification, were mainly found enriched in QJJ sediments, while dissimilatory nitrate reduction and assimilatory nitrate reduction modules were found enriched in DBC and QJJ plankton, respectively. For sulphur metabolism, thiosulfate oxidation was found to be abundant in DBC plankton, while dissimilatory sulphate reduction was found enriched in BLK sediments (Fig. 2d). These findings suggest that distinct differences in microbial metabolic pathways were found among these four lakes.

**Fig. 2.**
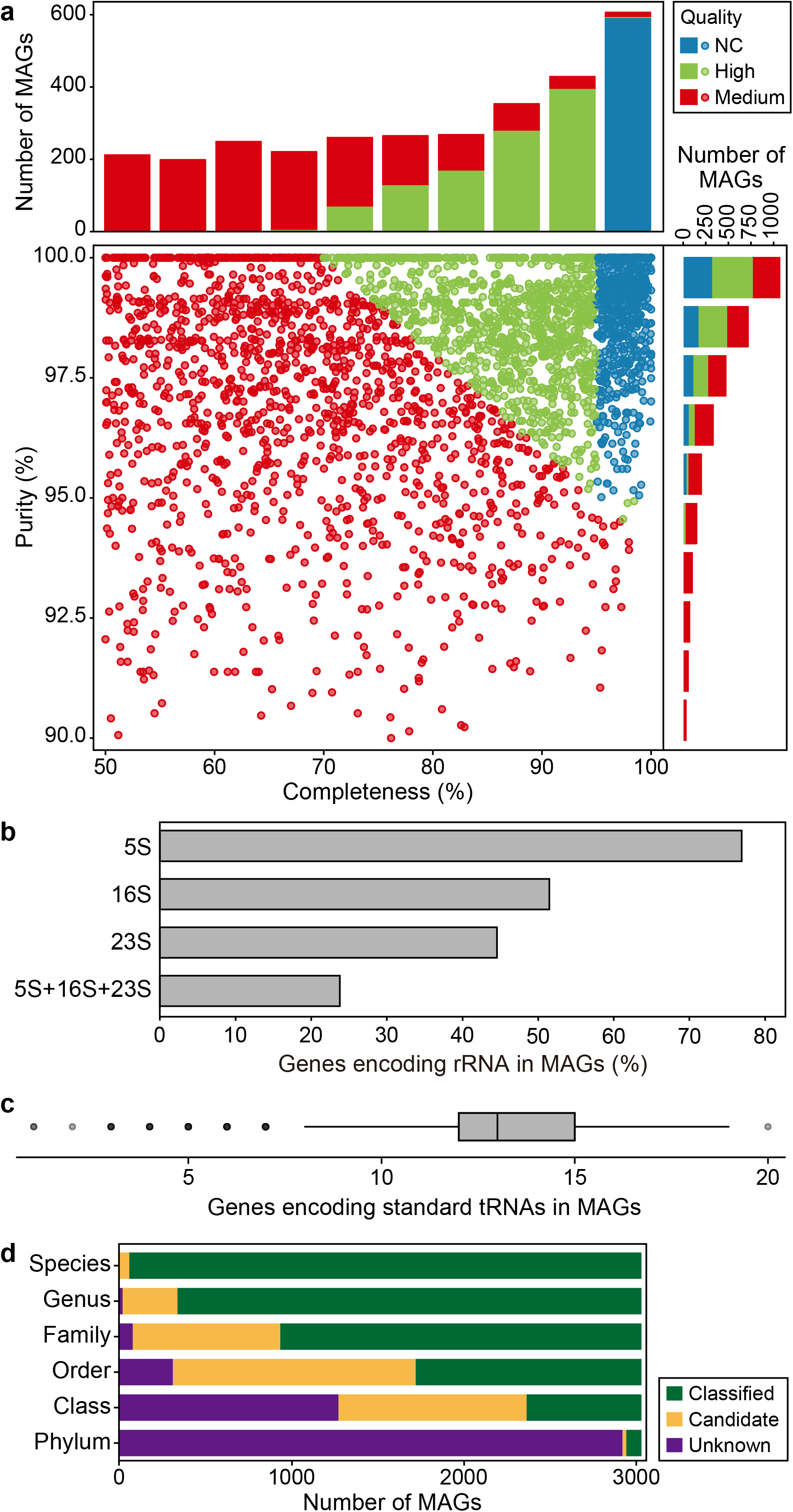
Microbial composition and genetic characteristics of four saline lakes. **a**, Alpha indices of predicted genes from lakewater and sedimentary communities in different lakes. Red: ADH (Aiding Lake), blue: BLK (Barkol Lake), green: DBC (Dabancheng Lake), purple: QJJ (Qijiaojing Lake). Asterisks indicate statistical significance (*P* < 0.05). NS: No statistical significance (*P* > 0.05). **b**, Distance-based Redundancy Analysis (dbRDA) of predicted genes from lakewater and sedimentary communities in different lakes. TOC: Total Organic Carbon, Li^+^: lithium, Na^+^: sodium, NH_4_^+^: ammonium, K^+^: potassium, Mg^2+^: magnesium, Ca^2+^: calcium, Cl^−^: chloridion, SO_4_^2−^: sulphate. **c**, Relative abundance of genes involved in key pathways, including carbon fixation, fatty acid metabolism, methane metabolism, nitrogen metabolism, photosynthesis, and sulfur metabolism. **d**, Z-score normalised relative abundance of modules in carbon fixation, methane, nitrogen, and sulfur metabolism.

**Fig. 3.**
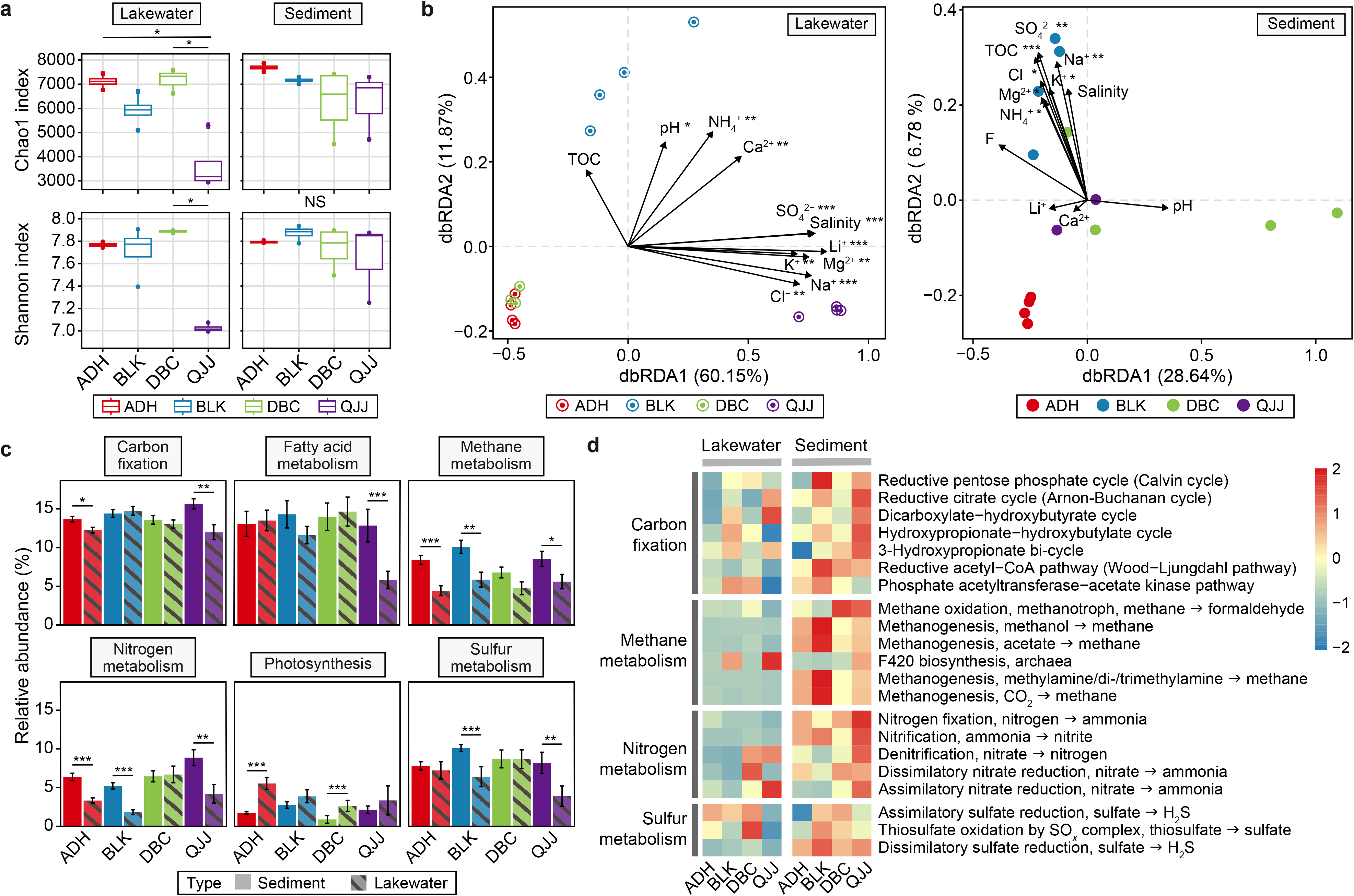
Phylogenetic analysis of MDM MAGs. Unrooted maximum likelihood trees of archaeal and bacterial MAGs were constructed with 122 and 120 concatenated marker genes in IQ-TREE using Q.pfam+R10 and LG+F+R10 models, respectively. **a**, MDM proportion of lakewater and sedimentary communities in terms of relative abundance and number of MAGs. Red: MDMs; Grey: non-MDMs. **b**, Microbial structure of lakewater and sedimentary communities in different lakes regarding relative abundance and number of MAGs classified at the phylum level. **c**,**d**, Phylogenetic tree of bacterial (**c**) and archaeal (**d**) MAGs. MDM lineages (bacterial) or phyla (archaeal) were highlighted in red, and MDM MAG(s) present in the corresponding lineages/phyla were indicated with shapes. ADH: square, BLK: circle, DBC: triangle, QJJ: diamond, solid symbol: sediment, hallowed symbol: plankton.

### 3.3. Microbial dark matters were highly diverse in the hypersaline lakes

Among the 3,030 non-redundant MAGs, 2,685 were classified as bacteria spanning 70 phyla, with 18 (25.7%) considered as putatively novel phyla. Here, a novel phylum was defined when the majority of microbial species within this phylum were not amenable to being isolated under standard laboratory techniques. The remaining 345 MAGs were identified as archaea, spanning 12 phyla, including eight putatively novel phyla. While a vast majority of MAGs were assigned to known phyla (96.6% in plankton and 92.5% in sediment in relative abundance), those minority MAGs in putatively novel phyla demonstrated higher diversity that comprised 16.7% and 30.1% of total MAGs in numbers in the plankton and sediments, respectively (Fig. 2a). This observation suggested that these putatively novel MAGs could exhibit considerable diversity in hypersaline environments. Lineages of common phyla were ubiquitously found across most samples, but distinct distributions of putatively novel phyla (except Patescibacteria) were only identified in specific lakes or sample types (Fig. 2b). These putatively novel phyla were predominantly located in samples with high microbial diversity (i.e., ADH and BLK sediment, Fig. 2b,c), but low relative abundance (<0.5%) among recovered MAGs, indicating that a considerable number of unknown microorganisms inhabitant in these environments were yet to be discovered.

### 3.4. Biosynthetic Potentials of putatively novel MAGs in saline lakes

In the 3,030 hypersaline lake MAGs, we identified 9,635 biosynthetic gene clusters (BGCs), including 232 BGCs clustered into known gene cluster families (GCFs) at 0.5 distance cutoff. A total of 9,403 BGCs (97.6% of overall BGCs) that were not clustered to MiBIG BGCs were described as unknown BGCs (Fig. 4a). Notably, several putatively novel bacterial phyla were enriched in BGCs (BGCs per MAG > 5), including OLB16, Moduliflexota, KSB1, Fibrobacterota, and B130-G9. This enrichment was only observed in Planctomycetota, Myxococcota, and Acidobacteriota among the known phyla (Fig. 4b), suggesting that putatively novel bacterial MAGs in these saline lakes may possess abundant biosynthetic potentials. Fewer BGCs were found in archaeal MAGs (BGCs per MAG < 2), possibly due to the insufficient data collected in the database.

**Fig. 4.**
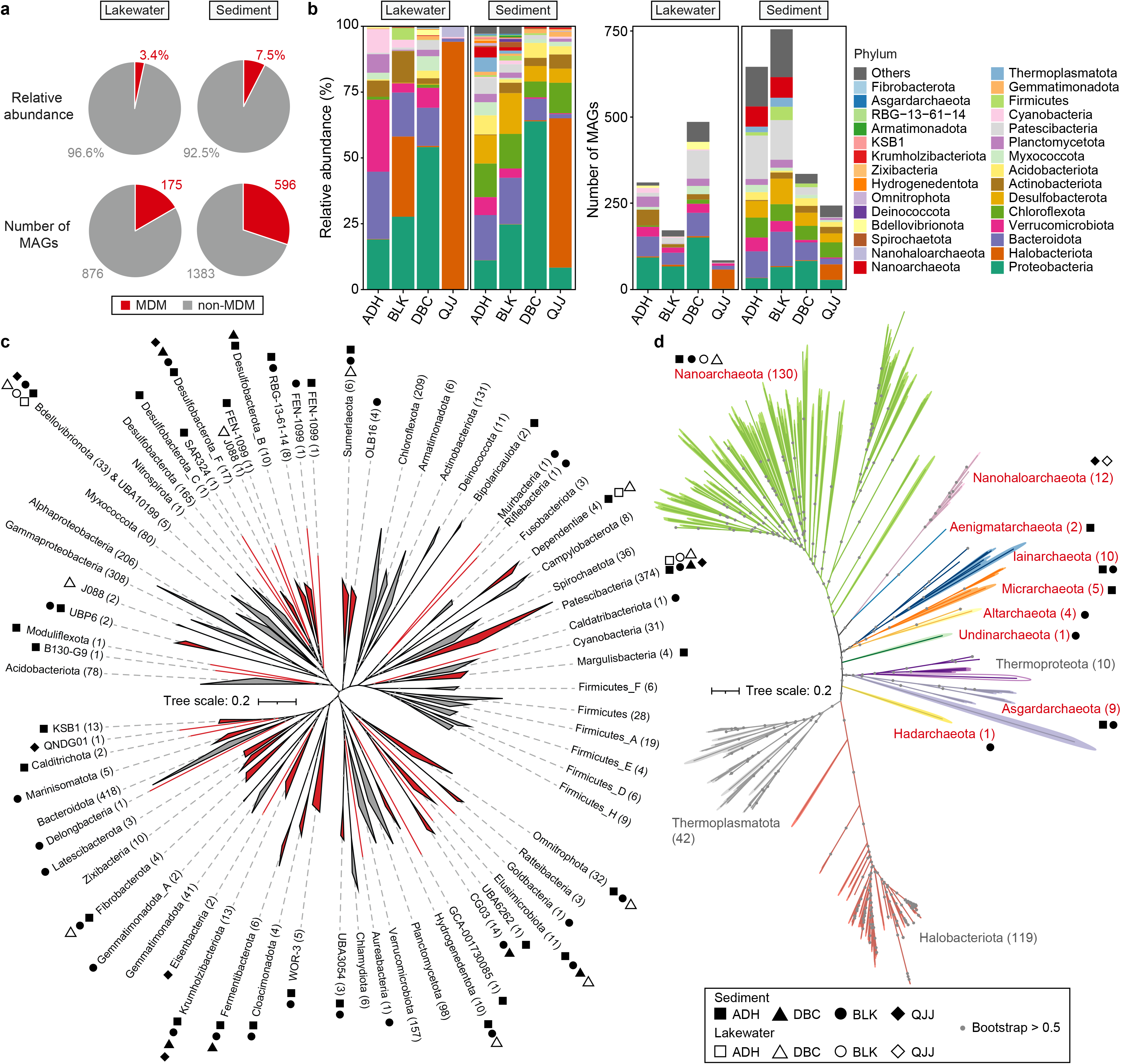
Analysis of biosynthetic potentials of hypersaline lake MAGs. **a**, A summary of predicted BGCs. In the pie chart on top, blue indicates that predicted BGCs were clustered to known BGCs in the MiBIG database; grey indicates that BGCs were not clustered to a known BGC in the MiBIG database. The pie chart at the bottom summarised the number of BGCs annotated to a BGC class. **b**, Normalised number of BGCs per MAG in each phylum. **c**, Phylogenetic trees of archaeal and bacterial MAGs with completeness ≥ 70, or number of BGCs ≥ 5. Midpoint-rooted maximum likelihood trees of archaeal and bacterial MAGs were constructed with 122 and 120 concatenated marker genes in IQ-TREE using Q.pfam+R10 and LG+F+R10 models, respectively. Black branches indicate bacterial MAGs, while purple branches indicate archaeal MAGs.

To further explore biosynthetic potentials of putatively novel MAGs, bacterial and archaeal phylogenetic trees were constructed, selecting MAGs with Completeness ≥ 70% or BGCs ≥ 5. Sixteen bacterial and two archaeal phyla contained MAG(s) with BGCs ≥ 5, suggesting potential biosynthetic enrichment in these phyla. Bacterial and archaeal phyla with small genome sizes (i.e., Patescibacteria and Nanoarchaeota) generally lacked BGCs, except one MAG from Nanoarchaeota, considered an outlier. Some phyla with only a single MAG obtained from saline lake metagenomes, such as B130-G9, Moduliflexota, UBP6, etc. (labelled individually on the tree, Fig. 4b), were not further discussed in this study as their biosynthetic pattern and consistency cannot be validated due to insufficient data. However, other putatively novel phyla consistently demonstrated enrichment with specific BGC Classes. For example, phylum Armatimonadota primarily consisted of RiPPs, Terpene and Other BGC classes, while phylum OLB16 was enriched in NRPS, RiPPs, and Terpene. Most of the MAGs in phylum Bdellovibrionota had BGCs in Terpene, RiPPs, and Other BGC classes, whereas KSB1 MAGs featured NRPS, PKS-NRPS hybrid, and Terpene classes (Fig. 4c). These observations suggest that MAGs within these phyla may have similar biosynthetic potentials. Two Asgardarchaeota MAGs were found to be enriched with BGCs, although this pattern was not consistent across the entire phyla, necessitating more genome analysis for further exploration.

### 3.5. Biodegradation potentials of putatively novel MAGs in saline lakes

To dissect the biodegradation potentials of hypersaline lake microorganisms, we analysed genes involved in the nitrogen cycle, dehalogenation, plastics degradation, and PAHs degradation within putatively novel MAGs. Several putatively novel bacterial phyla were found to be rich in nitrogen metabolism genes, such as GCA-001730085, Hydrogenedentota, OLB16, Fibrobacterota, Krumholzibacteriota, FEN-1099, RBG-13-61-14, and Zixibacteria (Fig. 5a), suggesting that these putatively novel MAGs may play a crucial role in the nitrogen cycle in hypersaline lakes. Interestingly, although MAGs belong to typical anaerobic ammonium oxidizing bacteria (AnAOB), i.e., families Scalinduaceae and Brocadiaceae in the order Brocadiales, class Brocadiae, were not found in this study, anaerobic ammonium oxidation (Anammox)-related genes (i.e., genes *hzs* and *hdh*) were found in several putatively novel phyla in addition to the known AnAOB groups, such as GCA-001730085, Hydrogenedentota, Calditrichota, KSB1, FEN-1099, and SAR324 (Fig. 5a). While fewer nitrogen cycle related genes were found in archaea than bacteria, we found that Asgard archaea universally contain *nirBD* and *norBC* genes, suggesting that they might be functionally involved in the nitrogen cycle in hypersaline lakes (Fig. 5b). In terms of the dehalogenation genes *rdh*, we found a number of putatively novel lineages with *rdh* genes in the hypersaline lake MAGs, such as bacterial phyla Eisenbacteria, Krumholzibacteriota, Zixibacteria, KSB1, and Marinisomatota (Fig. 5a). In archaeal taxa, *rdh* gene were found in classes Thorarchaeia, Lokiarchaeia, and phylum Hadarchaeota (Fig. 5b). These findings suggested that these putatively novel MAGs may have dehalogenation capacity in hypersaline lakes.

**Fig. 5.**
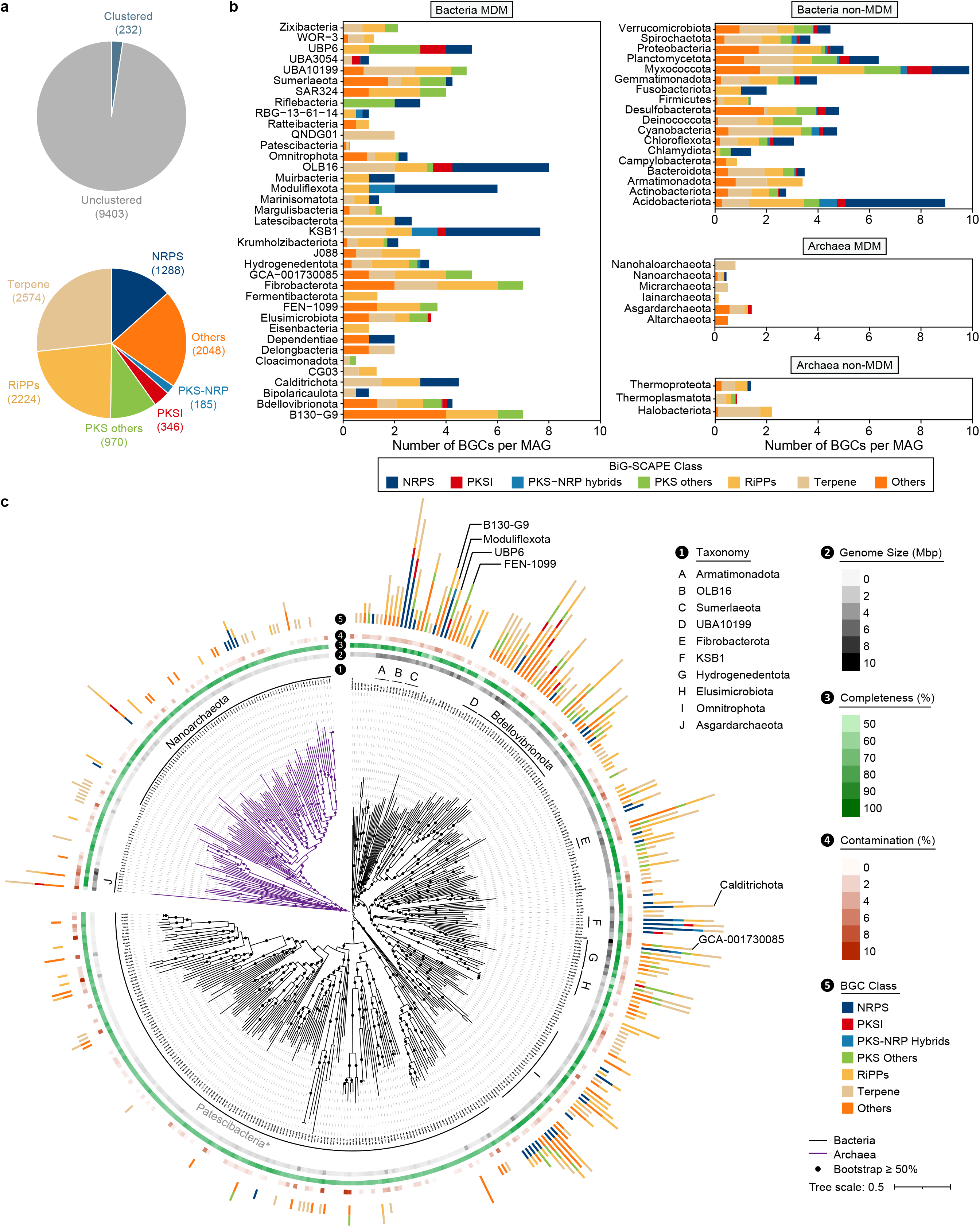
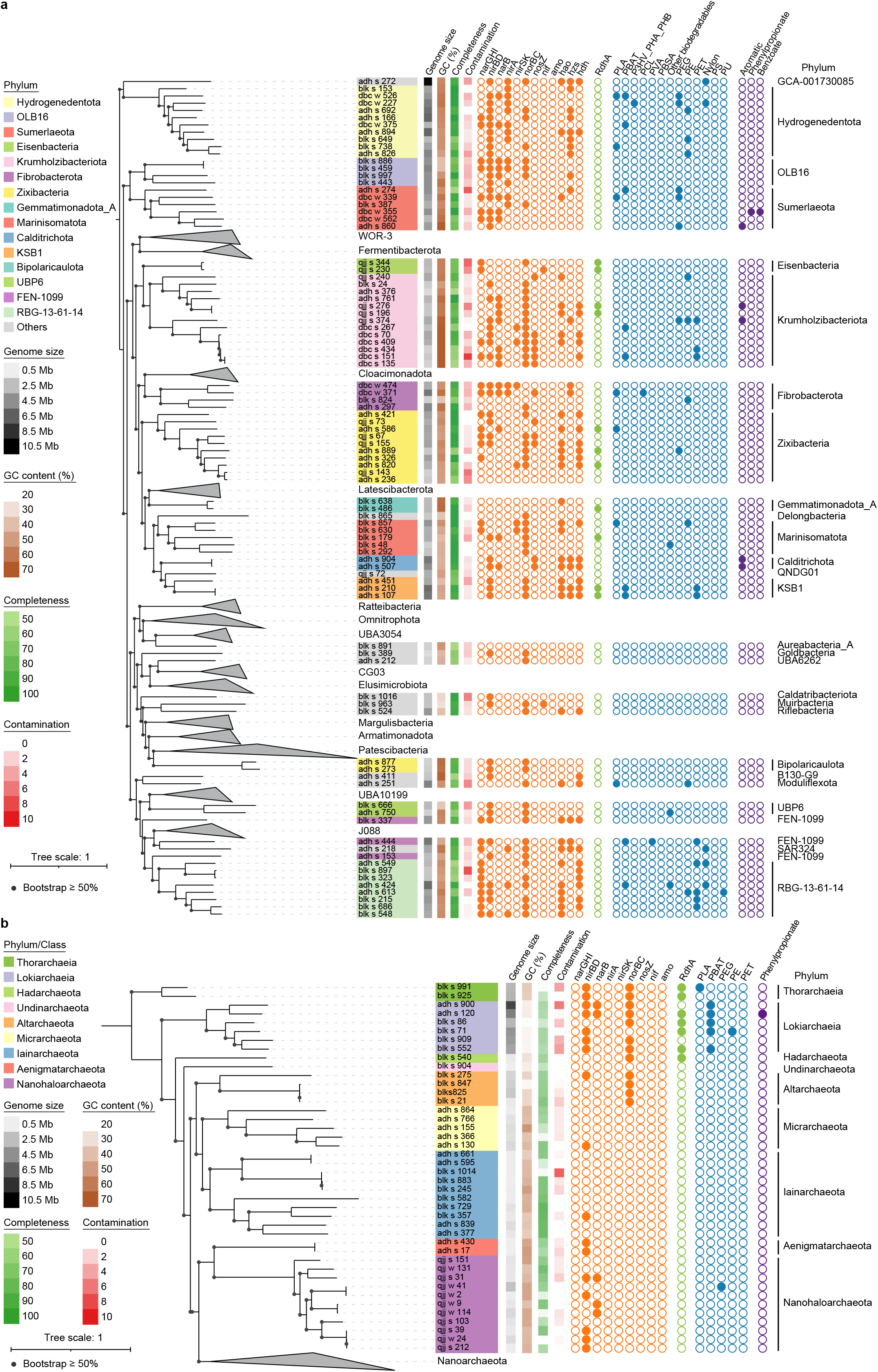
Phylogenetic analysis of biodegradative potentials in bacterial (**a**) and archaeal (**b**) MDM MAGs. Maximum likelihood trees of bacterial and archaeal MAGs were constructed with 120 and 122 concatenated marker genes in IQ-TREE using LG+F+R10 and Q.pfam+R10 models, respectively. The bacterial and archaeal trees were rooted with phylum GCA-001730085 and class Thorarchaeia, respectively. Key biodegradative genes were annotated in orange (nitrogen metabolism), green (dehalogenation), blue (plastics degradation), and purple (PAHs degradation).

To explore plastic degradation and PAHs potentials of saline lake MAGs, dereplicated ORFs were annotated against the PlasticDB and Darhd databases. Results showed that putatively novel MAGs with plastic degradation capabilities distributed in most of the phyla, including for previously known as non-biodegradable plastics, such as polyethylene (PE), polyethylene terephthalate (PET), polyamide (Nylon), polystyrene (PS), and polyurethane (PU), were found in phyla GCA-001730085, Hydrogenedentota, Krumholzibacteriota, Fibrobacterota, Marinisomatota, KSB1, Moduliflexota, FEN-1099, SAR324, RBG-13-61-14, and class Lokiarchaeia (Fig. 5a,b). For PAHs degradation potentials, genes such as aromatic ring-hydroxylating dioxygenase, phenylpropionate dioxygenase, and benzoate dioxygenase were found across putatively novel MAGs, spanning phyla Sumerlaeota, Krumholzibacteriota, Calditrichota, and class Lokiarchaeia (Fig. 5a,b), suggesting that many putatively novel MAGs may also have the potential to degrade microbially degradable plastics and PAHs.

## 4. Discussion

Uncharacterised microorganisms possess a vast repository of undiscovered enzymes and metabolic pathways that hold the potential for applications in various fields, including biotechnology, medicine, agriculture, and environmental remediation [49–54]. Hypersaline lakes are physiochemically diverse, leading to high complexity of microbial communities [55–59]. Thus, more microbial information with a broader range of genetic diversity could be unravelled from hypersaline lakes compared to engineered environments where specialised microbes were enriched [60–62]. In this study, we investigated four hypersaline lakes with distinct biochemical characteristics and successfully uncovered over 3,000 MAGs, providing valuable microbial resources at the genome-scale. Although salinity is typically found to have a negative correlation with microbial diversity [59, 63], halophilic archaea, such as Nanohaloarchaeota archaea, are uniquely present in some environments with extremely high salinity, such as QJJ lake (Fig. 3d). These archaea have been considered as symbionts, exhibiting intracellular lifestyles with other prokaryotic cells [24, 64]. Several putatively novel phyla were exclusively identified in ADH and BLK sediments (Fig. 3c,d). These putatively novel MAGs might correlate with TOC (in BLK) or other unknown environmental factors (in ADH), necessitating further exploration. Sulphate reduction bacteria (SRBs), such as Desulfobacterota and Zixibacteria, were abundant in BLK sediment, possibly due to the high concentration of SO_4_^2−^ and anaerobic conditions in the sediments. Furthermore, building upon our previous study on recovering Asgardarchaeota MAGs from ADH sediments [28], we have acquired new Asgardarchaeota MAGs from BLK sediments, which makes it the second where Asgardarchaeota MAGs have been recovered from deep-inland saline environments.

The recovery of 3,030 MAGs also provided valuable information on biosynthesis and biodegradation potentials. Although a metatranscriptomic analysis was not included in this study to confirm biosynthetic activity, our metagenomic analyses revealed nearly 10,000 potential BGCs, including ∼80% of which were considered novel. Notably, our findings concerning putatively novel phyla that are enriched in BGCs aligned with previous studies that included metatranscriptomic analysis from marine environments [65, 66], such as Omnitrophota, KSB1, OLB16, and FEN-1099, etc., further validated our findings in the biosynthetic analysis. On the other hand, biodegradation capabilities such as nitrogen removals (especially Anammox), dehalogenation, plastic degradation, and PAHs degradation are areas of intense interest for environmental remediation efforts [67–72]. The presence of biodegradation genes in putatively novel MAGs observed in this study offered new potentials for the degradation of pollutants, plastics, and toxic compounds. Biodegradation genes in a hypersaline lake environment may not be directly linked to microbial functionalities or serve as a bioindicator to plastics or PAHs pollution, given the absence of metatranscriptomic analyses. Nonetheless, our findings offered a genetic reservoir with the potentials for future biotechnological applications. Future studies should include metatranscriptomic analysis to confirm the expression of functional genes, and experimental validation is indispensable to substantiate specific microbial functions. Overall, these findings vastly improved our understanding of the putatively novel genomes in hypersaline lake environments, as well as the metabolic potentials of novel microbial species in developing biosynthetic and biodegradation capabilities, which can contribute to a broad range of applications, including biofuels, medicine, agricultural and industrial applications, as well as environmental remediation efforts. Moreover, the recovery of MAGs provided valuable genomic resources that enable insights into microbial evolution, potentially expanding our understanding of the Tree of Life [73].

Extreme environments are typically inhospitable to a large proportion of known microorganisms [17, 74]. However, such environments offer a group of specialised extremophiles that have evolved unique strategies to maintain their stability and functionality under such extreme conditions [75–77]. Recent studies have uncovered dozens of thousands of MAGs from various extreme environments, including deserts, oil fields, hydrothermal vents, cold seeps, hypersaline lakes, glaciers, etc. [15, 78–83]. Within putatively novel phyla, a higher diversity of Patescibacteria and Nanoarchaeota has been observed in hypersaline lake environments compared to other environments, aligning with previous findings of saline lake studies [15, 24]. Some MAGs, such as Caldatribacteriota and KSB1, were rarely reported in other environments except marine sediments [84, 85] or, as is the case of Margulisbacteria in underground water environments [86]. These observations may be possibly attributable to the low abundance of such microorganisms in specific environments. Microorganisms in hypersaline lakes are known as halotolerant or halophilic live forms, with distinct physiological and biochemical features that enable them to adapt to such extreme environments [87, 88]. Interestingly, while hypersaline lakes were previously known to be nutrient-poor, recent studies revealed that some of them are nutrient-rich, particularly in the lake sediment [89–91], which aligns with the results of this study. The remarkable adaptations and compatibilities of these organisms to high salinity environments, along with specialised functions including osmoregulation, “salt-in” strategy, membrane formation, and halophilic enzymes, etc. [92–96], offer exceptional opportunities for biotechnological applications. These applications range from non-sterile bioproduction and bioremediation of high salinity wastes to salt production and desalination. Collectively, mining and utilising microbial resources from hypersaline lakes can significantly advance our understanding of the metabolic potentials of extremophiles. This knowledge can be harnessed for applications in biotechnological industries and environmental management practices, broadening the spectrum of practical applications and contributing to sustainable development and environmental preservation.

Despite the promising potentials of exploiting uncultured hypersaline microorganisms (namely MDMs) for various applications, the limitations of exploiting these MDMs cannot be overlooked. One key challenge is that MDMs often contain undocumented genes or enzymes in the current databases. This can affect the accuracy of predictions, annotations, or classification in genomic analyses [49, 97, 98]. While advancements in state-of-art sequencing technologies and bioinformatic tools, such as long-read sequencing [99, 100], single-cell sequencing [101], deep refinement [28], and machine learning [102] approaches, have assisted in mitigating these discrepancies, each of these technologies has its limitations. These still hampers the robust *de novo* assembly of low abundant putatively novel MAGs from the microbial communities. Moreover, since successfully culturing a microorganism is still considered the “gold standard” approach of microbial characterisation, it is still challenging to replicate extreme conditions such as hypersaline environments under laboratory conditions, making it difficult to culture putatively novel MAGs. Even if the culturing approaches can be managed, validating their functional and metabolic properties to confirm biosynthesis and biodegradation potentials remains a major challenge. Cutting-edge technologies and in-depth analysis tools, such as multi-omics approaches [103], Raman spectroscopy methods [104], and high-throughput single-cell sequencing technologies [105], may assist in the isolation and culturing of uncharacterised microorganisms. However, most of these techniques have been sophisticatedly applied to human gut microbiome studies; thus, such application to extreme environmental samples remains challenging, both in terms of laboratory experiments and *in silico* analyses.

Based on the current findings of biodegradation potentials in this study, future research is proposed to be conducted on targeted culture enrichment. For example, potential Anammox bacteria could be enriched from specific lakes where MAGs with Anammox signature genes were found. To achieve this, multi-omics analysis approaches could be applied to assist the enrichment to reveal the nutrient requirement of target species, expression of key genes, and interactions with other microorganisms in the enrichment systems [106, 107]. However, further exploration of putatively novel microorganisms from hypersaline lakes and other extreme environments is complex, with numerous technical challenges to be addressed. One of the key conundrums lies in the increase of resolution for low coverage of microbial genomes, which is difficult to recover via binning. The successful enrichment of these putatively novel microorganisms is also hindered by intricate nutrient requirements, complex microbial interactions, and slow growth rates associated with these extremophiles [108–110]. Possible solutions might be conceived such as (1) the utilisation of advanced sequencing technologies, such as Hi-C, Pore-C, and HiFi sequencing [111–113], which could potentially yield higher-quality and more complete genomic assemblies from metagenomic datasets, offering a more detailed and accurate view of the landscape of putatively novel genomes; (2) the development of new bioinformatic tools to enhance the accuracy and resolution of recovered genomes from metagenomic datasets, potentially achieved through the application of machine learning algorithms or the improvement of existing bioinformatic tools; (3) the upgraded laboratory techniques for culture enrichment (e.g., machine learning-based high-throughput microbial culturomics) [114] or “counterselection” strategy on extreme environmental metagenomes. Nevertheless, successfully developing and applying these approaches could significantly advance our discovery of MDMs and their associated biotechnological potentials. It could also unfold possibilities for exploration of life in the most extreme environments on Earth, including abyssal, hadal, deep subsurface, and even beyond, into extraterrestrial environments.

## 5. Conclusions

Our study implemented genome-centric analyses on over 3,000 MAGs recovered from four hypersaline lakes in Xinjiang Uygur Zizhiqu, China, each characterised by distinct environmental parameters. This effort has significantly augmented the repository of genomic data and valuable insights into microbial life in extreme environments, particularly hypersaline lakes. We found over 8,000 potential BGCs and uncovered several putatively novel phyla that may be enriched in biosynthetic capacities, substantially supporting future research in biotechnological applications. In addition, the discovery of biodegradation genes within certain putatively novel lineages also suggests promising prospects for environmental remediation strategies. In summary, this study expands our knowledge of microbial diversity and function in extreme environments, paving the path to the future discovery of uncultured microorganisms. This research deepens our understanding of the Tree of Life and unveils avenues for diverse applications in biotechnology and environmental remediation.

## CRediT authorship contribution statement

**Zhiguang Qiu:** Conceptualisation, Methodology, Formal analysis, Writing – Original Draft, Writing – Review & Editing, Visualisation, Validation. **Yuanyuan Zhu:** Formal analysis, Visualisation. **Qing Zhang:** Formal analysis. **Xuejiao Qiao:** Conceptualisation, Resources. **Rong Mu:** Investigation. **Zheng Xu:** Conceptualisation, Visualisation. **Yan Yan:** Resources. **Fan Wang:** Resources. **Tong Zhang:** Writing – Review & Editing. **Wei-Qin Zhuang:** Writing – Review & Editing. **Ke Yu:** Conceptualisation, Resources, Project administration, Funding acquisition.

## Acknowledgements

This research was supported by the National Key Research and Development Program of China (2021YFA1301300), Nature Science Foundation of China (62202014 and 61972217), Shenzhen Basic Research Programs (JCYJ20190808183205731, JCYJ20220812103301001, and JCYJ20220813151736001), the Shenzhen Knowledge Innovation Program Basic Research Project (JCYJ20190808183205731), and Science and Technology Planning Project of Shenzhen Municipality (JCYJ20200109120416654). The authors would like to thank Dr. Luhua Xie, Dr. Bing Li, Ms. Liyu Zhang, and Ms. Xiaoling Gao for their technical assistance and valuable discussion.

## Availability of data and materials

Metagenomic sequence data of saline lakes are available in the CNCB-NGDC database under project PRJCA014712.

---

**Fig. S1.** Location of Aiding Lake (ADH), Barkol Lake (BLK), Dabancheng Lake (DBC), and Qijiaojing Lake (QJJ).

**Fig. S2.** NMDS plot of genetic diversities analysed in Bray-Curtis dissimilarities. Different colours indicate different lakes, including red (ADH), cyan (BLK), green (DBC), and purple (QJJ), while different shapes indicate different lake compartments, including sediment (solid circle) and plankton (hollowed circle).

## References

[1] M. Bahram, F. Hildebrand, S. K. Forslund, J. L. Anderson, N. A. Soudzilovskaia, P. M. Bodegom, J. Bengtsson-Palme, S. Anslan, L. P. Coelho and H. Harend. Structure and function of the global topsoil microbiome. Nature 560(7717) (2018), 233–237.

[2] P. G. Falkowski, T. Fenchel and E. F. Delong. The microbial engines that drive Earth’s biogeochemical cycles. science 320(5879) (2008), 1034–1039.

[3] S. Louca, L. W. Parfrey and M. Doebeli. Decoupling function and taxonomy in the global ocean microbiome. Science 353(6305) (2016), 1272–1277.

[4] R. Sender, S. Fuchs and R. Milo. Revised estimates for the number of human and bacteria cells in the body. PLoS Biol. 14(8) (2016), e1002533.

[5] H. Barabadi, B. Tajani, M. Moradi, K. Damavandi Kamali, R. Meena, S. Honary, M. A. Mahjoub and M. Saravanan. Penicillium family as emerging nanofactory for biosynthesis of green nanomaterials: a journey into the world of microorganisms. Journal of Cluster Science 30 (2019), 843–856.

[6] J. A. Chemler and M. A. Koffas. Metabolic engineering for plant natural product biosynthesis in microbes. Curr. Opin. Biotechnol. 19(6) (2008), 597–605.

[7] J. Zhang, Y. Peng, X. Li and R. Du. Feasibility of partial-denitrification/anammox for pharmaceutical wastewater treatment in a hybrid biofilm reactor. Water Res. 208 (2022), 117856.

[8] S. Wang, K. R. Chng, A. Wilm, S. Zhao, K.-L. Yang, N. Nagarajan and J. He. Genomic characterization of three unique Dehalococcoides that respire on persistent polychlorinated biphenyls. Proceedings of the National Academy of Sciences 111(33) (2014), 12103–12108.

[9] R. Zallot, N. Oberg and J. A. Gerlt. Discovery of new enzymatic functions and metabolic pathways using genomic enzymology web tools. Curr. Opin. Biotechnol. 69 (2021), 77–90.

[10] A. Saravanan, P. S. Kumar, S. Karishma, D.-V. N. Vo, S. Jeevanantham, P. Yaashikaa and C. S. George. A review on biosynthesis of metal nanoparticles and its environmental applications. Chemosphere 264 (2021), 128580.

[11] J. Wu, G. Du, J. Zhou and J. Chen. Systems metabolic engineering of microorganisms to achieve large-scale production of flavonoid scaffolds. J. Biotechnol. 188 (2014), 72–80.

[12] N. Taghavi, I. A. Udugama, W.-Q. Zhuang and S. Baroutian. Challenges in biodegradation of non-degradable thermoplastic waste: From environmental impact to operational readiness. Biotechnol. Adv. 49 (2021), 107731.

[13] C. C. Azubuike, C. B. Chikere and G. C. Okpokwasili. Bioremediation techniques–classification based on site of application: principles, advantages, limitations and prospects. World Journal of Microbiology and Biotechnology 32 (2016), 1–18.

[14] A. Ventosa, R. R. de la Haba, C. Sanchez-Porro and R. T. Papke. Microbial diversity of hypersaline environments: a metagenomic approach. Curr. Opin. Microbiol. 25 (2015), 80–87.

[15] C. D. Vavourakis, A.-S. Andrei, M. Mehrshad, R. Ghai, D. Y. Sorokin and G. Muyzer. A metagenomics roadmap to the uncultured genome diversity in hypersaline soda lake sediments. Microbiome 6(1) (2018), 1–18.

[16] D. Y. Sorokin, T. Berben, E. D. Melton, L. Overmars, C. D. Vavourakis and G. Muyzer. Microbial diversity and biogeochemical cycling in soda lakes. Extremophiles 18 (2014), 791–809.

[17] L. J. Rothschild and R. L. Mancinelli. Life in extreme environments. Nature 409(6823) (2001), 1092–1101.

[18] W.-S. Shu and L.-N. Huang. Microbial diversity in extreme environments. Nature Reviews Microbiology 20(4) (2022), 219–235.

[19] L. Krienitz and K. Kotut. Fluctuating algal food populations and the occurrence of lesser flamingos (Phoeniconaias minor) in three Kenyan rift valley lakes 1. J. Phycol. 46(6) (2010), 1088–1096.

[20] P. Rojas, N. Rodríguez, V. de la Fuente, D. Sánchez-Mata, R. Amils and J. L. Sanz. Microbial diversity associated with the anaerobic sediments of a soda lake (Mono Lake, California, USA). Canadian journal of microbiology 64(6) (2018), 385–392.

[21] N. M. Mesbah and J. Wiegel. Life under multiple extreme conditions: diversity and physiology of the halophilic alkalithermophiles. Appl. Environ. Microbiol. 78(12) (2012), 4074–4082.

[22] J. Schultz and A. S. Rosado. Extreme environments: a source of biosurfactants for biotechnological applications. Extremophiles 24 (2020), 189–206.

[23] D. Zhao, S. Zhang, Q. Xue, J. Chen, J. Zhou, F. Cheng, M. Li, Y. Zhu, H. Yu and S. Hu. Abundant taxa and favorable pathways in the microbiome of soda-saline lakes in Inner Mongolia. Frontiers in microbiology 11 (2020), 1740.

[24] Y.-G. Xie, Z.-H. Luo, B.-Z. Fang, J.-Y. Jiao, Q.-J. Xie, X.-R. Cao, Y.-N. Qu, Y.-L. Qi, Y.-Z. Rao and Y.-X. Li. Functional differentiation determines the molecular basis of the symbiotic lifestyle of Ca. Nanohaloarchaeota. Microbiome 10(1) (2022), 1–13.

[25] M. Ji, W. Kong, L. Yue, J. Wang, Y. Deng and L. Zhu. Salinity reduces bacterial diversity, but increases network complexity in Tibetan Plateau lakes. FEMS Microbiol. Ecol. 95(12) (2019), fiz190.

[26] S. Chen, Y. Zhou, Y. Chen and J. Gu. fastp: an ultra-fast all-in-one FASTQ preprocessor. Bioinformatics 34(17) (2018), i884–i890.

[27] A. Bankevich, S. Nurk, D. Antipov, A. A. Gurevich, M. Dvorkin, A. S. Kulikov, V. M. Lesin, S. I. Nikolenko, S. Pham and A. D. Prjibelski. SPAdes: a new genome assembly algorithm and its applications to single-cell sequencing. J. Comput. Biol. 19(5) (2012), 455–477.

[28] K. Yu, Z. Qiu, R. Mu, X. Qiao, L. Zhang, C.-A. Lian, C. Deng, Y. Wu, Z. Xu, B. Li, B. Pan, Y. Zhang, L. Fan, Y.-x. Liu, H. Cao, T. Jin, B. Chen, F. Wang, Y. Yan, L. Xie, L. Zhou, S. Yi, S. Chi, C. Zhang, T. Zhang and W. Zhuang. Recovery of high-qualitied Genomes from a deep-inland Salt Lake Using BASALT. bioRxiv (2021), 2021.2003.2005.434042.

[29] D. H. Parks, M. Imelfort, C. T. Skennerton, P. Hugenholtz and G. W. Tyson. CheckM: assessing the quality of microbial genomes recovered from isolates, single cells, and metagenomes. Genome Res. 25(7) (2015), 1043–1055.

[30] R. M. Bowers, N. C. Kyrpides, R. Stepanauskas, M. Harmon-Smith, D. Doud, T. Reddy, F. Schulz, J. Jarett, A. R. Rivers and E. A. Eloe-Fadrosh. Minimum information about a single amplified genome (MISAG) and a metagenome-assembled genome (MIMAG) of bacteria and archaea. Nat. Biotechnol. 35(8) (2017), 725–731.

[31] P.-A. Chaumeil, A. J. Mussig, P. Hugenholtz and D. H. Parks. GTDB-Tk: a toolkit to classify genomes with the Genome Taxonomy Database. Bioinformatics (2020).

[32] S. R. Eddy. Accelerated profile HMM searches. PLoS Comp. Biol. 7(10) (2011), e1002195.

[33] B. Q. Minh, H. A. Schmidt, O. Chernomor, D. Schrempf, M. D. Woodhams, A. Von Haeseler and R. Lanfear. IQ-TREE 2: new models and efficient methods for phylogenetic inference in the genomic era. Mol. Biol. Evol. 37(5) (2020), 1530–1534.

[34] I. Letunic and P. Bork. Interactive Tree Of Life (iTOL) v5: an online tool for phylogenetic tree display and annotation. Nucleic Acids Res. 49(W1) (2021), W293–W296.

[35] P. P. Chan, B. Y. Lin, A. J. Mak and T. M. Lowe. tRNAscan-SE 2.0: improved detection and functional classification of transfer RNA genes. Nucleic Acids Res. 49(16) (2021), 9077–9096.

[36] D. Hyatt, G.-L. Chen, P. F. LoCascio, M. L. Land, F. W. Larimer and L. J. Hauser. Prodigal: prokaryotic gene recognition and translation initiation site identification. Bioinformatics 11(1) (2010), 119.

[37] W. Li and A. Godzik. Cd-hit: a fast program for clustering and comparing large sets of protein or nucleotide sequences. Bioinformatics 22 (2006).

[38] B. Buchfink, C. Xie and D. H. Huson. Fast and sensitive protein alignment using DIAMOND. Nature Methods 12(1) (2015), 59.

[39] P. Jones, D. Binns, H.-Y. Chang, M. Fraser, W. Li, C. McAnulla, H. McWilliam, J. Maslen, A. Mitchell and G. Nuka. InterProScan 5: genome-scale protein function classification. Bioinformatics 30(9) (2014), 1236–1240.

[40] W. B. Langdon. Performance of genetic programming optimised Bowtie2 on genome comparison and analytic testing (GCAT) benchmarks. BioData mining 8(1) (2015), 1–7.

[41] H. Li, B. Handsaker, A. Wysoker, T. Fennell, J. Ruan, N. Homer, G. Marth, G. Abecasis and R. Durbin. The sequence alignment/map format and SAMtools. Bioinformatics 25 (2009).

[42] M. Albertsen, P. Hugenholtz, A. Skarshewski, K. L. Nielsen, G. W. Tyson and P. H. Nielsen. Genome sequences of rare, uncultured bacteria obtained by differential coverage binning of multiple metagenomes. Nat. Biotechnol. 31(6) (2013), 533–538.

[43] G. M. Douglas, V. J. Maffei, J. R. Zaneveld, S. N. Yurgel, J. R. Brown, C. M. Taylor, C. Huttenhower and M. G. I. Langille. PICRUSt2 for prediction of metagenome functions. Nat. Biotechnol. 38(6) (2020), 685–688.

[44] K. Blin, S. Shaw, A. M. Kloosterman, Z. Charlop-Powers, G. P. Van Wezel, M. H. Medema and T. Weber. antiSMASH 6.0: improving cluster detection and comparison capabilities. Nucleic Acids Res. 49(W1) (2021), W29–W35.

[45] J. C. Navarro-Muñoz, N. Selem-Mojica, M. W. Mullowney, S. A. Kautsar, J. H. Tryon, E. I. Parkinson, E. L. De Los Santos, M. Yeong, P. Cruz-Morales and S. Abubucker. A computational framework to explore large-scale biosynthetic diversity. Nat. Chem. Biol. 16(1) (2020), 60–68.

[46] S. A. Kautsar, K. Blin, S. Shaw, J. C. Navarro-Muñoz, B. R. Terlouw, J. J. Van Der Hooft, J. A. Van Santen, V. Tracanna, H. G. Suarez Duran and V. Pascal Andreu. MIBiG 2.0: a repository for biosynthetic gene clusters of known function. Nucleic Acids Res. 48(D1) (2020), D454–D458.

[47] S. Li, W. Shen, S. Lian, Y. Wu, Y. Qu and Y. Deng. Darhd: A sequence database for aromatic ring-hydroxylating dioxygenase analysis and primer evaluation. J. Hazard. Mater. 436 (2022), 129230.

[48] V. Gambarini, O. Pantos, J. M. Kingsbury, L. Weaver, K. M. Handley and G. Lear. PlasticDB: a database of microorganisms and proteins linked to plastic biodegradation. Database 2022 (2022).

[49] C. Rinke, P. Schwientek, A. Sczyrba, N. N. Ivanova, I. J. Anderson, J.-F. Cheng, A. Darling, S. Malfatti, B. K. Swan and E. A. Gies. Insights into the phylogeny and coding potential of microbial dark matter. Nature 499(7459) (2013), 431–437.

[50] A. Bhushan, P. J. Egli, E. E. Peters, M. F. Freeman and J. Piel. Genome mining-and synthetic biology-enabled production of hypermodified peptides. Nature chemistry 11(10) (2019), 931–939.

[51] V. Waschulin, C. Borsetto, R. James, K. K. Newsham, S. Donadio, C. Corre and E. Wellington. Biosynthetic potential of uncultured Antarctic soil bacteria revealed through long-read metagenomic sequencing. The ISME journal 16(1) (2022), 101–111.

[52] A. G. Atanasov, S. B. Zotchev, V. M. Dirsch and C. T. Supuran. Natural products in drug discovery: advances and opportunities. Nature reviews Drug discovery 20(3) (2021), 200–216.

[53] Z. Xu, T.-J. Park and H. Cao. Advances in mining and expressing microbial biosynthetic gene clusters. Crit. Rev. Microbiol. (2022), 1–20.

[54] S.-C. Chen, R. Budhraja, L. Adrian, F. Calabrese, H. Stryhanyuk, N. Musat, H.-H. Richnow, G.-L. Duan, Y.-G. Zhu and F. Musat. Novel clades of soil biphenyl degraders revealed by integrating isotope probing, multi-omics, and single-cell analyses. The ISME Journal 15(12) (2021), 3508–3521.

[55] R. Han, X. Zhang, J. Liu, Q. Long, L. Chen, D. Liu and D. Zhu. Microbial community structure and diversity within hypersaline Keke Salt Lake environments. Canadian journal of microbiology 63(11) (2017), 895–908.

[56] A.-Ş. Andrei, M. S. Robeson, A. Baricz, C. Coman, V. Muntean, A. Ionescu, G. Etiope, M. Alexe, C. I. Sicora and M. Podar. Contrasting taxonomic stratification of microbial communities in two hypersaline meromictic lakes. The ISME journal 9(12) (2015), 2642–2656.

[57] A. Baricz, C. M. Chiriac, A. Ș. Andrei, P. A. Bulzu, E. A. Levei, O. Cadar, K. P. Battes, M. Cîmpean, M. Șenilă and A. Cristea. Spatio-temporal insights into microbiology of the freshwater-to-hypersaline, oxic-hypoxic-euxinic waters of Ursu Lake. Environ. Microbiol. 23(7) (2021), 3523–3540.

[58] Y. Zhang, K. Yang, H. Chen, Y. Dong and W. Li. Origin, composition, and accumulation of dissolved organic matter in a hypersaline lake of the Qinghai-Tibet Plateau. Sci. Total Environ. 868 (2023), 161612.

[59] L. Wang, C. Lian, W. Wan, Z. Qiu, X. Luo, Q. Huang, Y. Deng, T. Zhang and K. Yu. Salinity-triggered homogeneous selection constrains the microbial function and stability in lakes. Appl. Microbiol. Biotechnol. (2023), 1–15.

[60] M. Naufal, J.-H. Wu and Y.-H. Shao. Glutamate enhances osmoadaptation of anammox bacteria under high salinity: genomic analysis and experimental evidence. Environ. Sci. Technol. 56(16) (2022), 11310–11322.

[61] K. Hu, D. Xu and Y. Chen. An assessment of sulfate reducing bacteria on treating sulfate-rich metal-laden wastewater from electroplating plant. J. Hazard. Mater. 393 (2020), 122376.

[62] H. A. Oyewusi, R. A. Wahab, Y. Kaya, M. F. Edbeib and F. Huyop. Alternative bioremediation agents against haloacids, haloacetates and chlorpyrifos using novel halogen-degrading bacterial isolates from the hypersaline lake Tuz. Catalysts 10(6) (2020), 651.

[63] J. Yang, L. a. Ma, H. Jiang, G. Wu and H. Dong. Salinity shapes microbial diversity and community structure in surface sediments of the Qinghai-Tibetan Lakes. Scientific reports 6(1) (2016), 25078.

[64] J. Hamm, Y. Liao, A. von Kügelgen, N. Dombrowski, E. Landers, C. Brownlee, E. Johansson, R. Whan, M. Baker and B. Baum. The intracellular lifestyle of an archaeal symbiont. (2023).

[65] D. Geller-McGrath, P. Mara, G. T. Taylor, E. Suter, V. Edgcomb and M. Pachiadaki. Diverse secondary metabolites are expressed in particle-associated and free-living microorganisms of the permanently anoxic Cariaco Basin. Nature Communications 14(1) (2023), 656.

[66] L. Paoli, H.-J. Ruscheweyh, C. C. Forneris, F. Hubrich, S. Kautsar, A. Bhushan, A. Lotti, Q. Clayssen, G. Salazar and A. Milanese. Biosynthetic potential of the global ocean microbiome. Nature 607(7917) (2022), 111–118.

[67] L. Liu, Y. Wang, Y. Che, Y. Chen, Y. Xia, R. Luo, S. H. Cheng, C. Zheng and T. Zhang. High-quality bacterial genomes of a partial-nitritation/anammox system by an iterative hybrid assembly method. Microbiome 8 (2020), 1–17.

[68] X. Qiao, C. Fu, Y. Chen, F. Fang, Y. Zhang, L. Ding, K. Yang, B. Pan, N. Xu and K. Yu. Molecular insights into enhanced nitrogen removal induced by trace fluoroquinolone antibiotics in an anammox system. Bioresour. Technol. 374 (2023), 128784.

[69] H. He, Y. Li, R. Shen, H. Shim, Y. Zeng, S. Zhao, Q. Lu, B. Mai and S. Wang. Environmental occurrence and remediation of emerging organohalides: A review. Environ. Pollut. 290 (2021), 118060.

[70] J. Liu, G. Xu, S. Zhao, C. Chen, M. J. Rogers and J. He. Mechanistic and microbial ecological insights into the impacts of micro-and nano-plastics on microbial reductive dehalogenation of organohalide pollutants. J. Hazard. Mater. 448 (2023), 130895.

[71] S. Zhang, Z. Hu and H. Wang. Metagenomic analysis exhibited the co-metabolism of polycyclic aromatic hydrocarbons by bacterial community from estuarine sediment. Environ. Int. 129 (2019), 308–319.

[72] J. Zrimec, M. Kokina, S. Jonasson, F. Zorrilla and A. Zelezniak. Plastic-degrading potential across the global microbiome correlates with recent pollution trends. MBio 12(5) (2021), e02155–02121.

[73] L. A. Hug, B. J. Baker, K. Anantharaman, C. T. Brown, A. J. Probst, C. J. Castelle, C. N. Butterfield, A. W. Hernsdorf, Y. Amano and K. Ise. A new view of the tree of life. Nature microbiology 1(5) (2016), 1–6.

[74] Y. S. Lee and W. Park. Current challenges and future directions for bacterial self-healing concrete. Appl. Microbiol. Biotechnol. 102 (2018), 3059–3070.

[75] V. Malavasi, S. Soru and G. Cao. Extremophile microalgae: the potential for biotechnological application. J. Phycol. 56(3) (2020), 559–573.

[76] W. Yin, Y. Wang, L. Liu and J. He. Biofilms: the microbial “protective clothing” in extreme environments. International journal of molecular sciences 20(14) (2019), 3423.

[77] S. Wang, J. Wang, Z. Liu and B. Zhang. Unraveling diverse survival strategies of microorganisms to vanadium stress in aquatic environments. Water Res. 221 (2022), 118813.

[78] K. M. Finstad, A. J. Probst, B. C. Thomas, G. L. Andersen, C. Demergasso, A. Echeverría, R. G. Amundson and J. F. Banfield. Microbial community structure and the persistence of cyanobacterial populations in salt crusts of the hyperarid Atacama Desert from genome-resolved metagenomics. Frontiers in microbiology 8 (2017), 1435.

[79] M. O. Eze, S. A. Lütgert, H. Neubauer, A. Balouri, A. A. Kraft, A. Sieven, R. Daniel and B. Wemheuer. Metagenome assembly and metagenome-assembled genome sequences from a historical oil field located in Wietze, Germany. Microbiology Resource Announcements 9(21) (2020).

[80] R. E. Anderson, J. Reveillaud, E. Reddington, T. O. Delmont, A. M. Eren, J. M. McDermott, J. S. Seewald and J. A. Huber. Genomic variation in microbial populations inhabiting the marine subseafloor at deep-sea hydrothermal vents. Nature communications 8(1) (2017), 1–11.

[81] X. Dong, C. Zhang, Y. Peng, H.-X. Zhang, L.-D. Shi, G. Wei, C. R. Hubert, Y. Wang and C. Greening. Phylogenetically and catabolically diverse diazotrophs reside in deep-sea cold seep sediments. Nature Communications 13(1) (2022), 4885.

[82] J. Liu, Y. Zheng, H. Lin, X. Wang, M. Li, Y. Liu, M. Yu, M. Zhao, N. Pedentchouk and D. J. Lea-Smith. Proliferation of hydrocarbon-degrading microbes at the bottom of the Mariana Trench. Microbiome 7(1) (2019), 1–13.

[83] Y. Liu, M. Ji, T. Yu, J. Zaugg, A. M. Anesio, Z. Zhang, S. Hu, P. Hugenholtz, K. Liu and P. Liu. A genome and gene catalog of glacier microbiomes. Nat. Biotechnol. 40(9) (2022), 1341–1348.

[84] Q. Li, Y. Zhou, R. Lu, P. Zheng and Y. Wang. Phylogeny, distribution and potential metabolism of candidate bacterial phylum KSB1. PeerJ 10 (2022), e13241.

[85] T. R. Iasakov, T. A. Kanapatskiy, S. V. Toshchakov, A. A. Korzhenkov, M. O. Ulyanova and N. V. Pimenov. The Baltic Sea methane pockmark microbiome: The new insights into the patterns of relative abundance and ANME niche separation. Mar. Environ. Res. 173 (2022), 105533.

[86] P. B. Matheus Carnevali, F. Schulz, C. J. Castelle, R. S. Kantor, P. M. Shih, I. Sharon, J. M. Santini, M. R. Olm, Y. Amano and B. C. Thomas. Hydrogen-based metabolism as an ancestral trait in lineages sibling to the Cyanobacteria. Nature communications 10(1) (2019), 463.

[87] D.-S. Mu, S. Wang, Q.-Y. Liang, Z.-Z. Du, R. Tian, Y. Ouyang, X.-P. Wang, A. Zhou, Y. Gong and G.-J. Chen. Bradymonabacteria, a novel bacterial predator group with versatile survival strategies in saline environments. Microbiome 8 (2020), 1–15.

[88] Y.-H. Liu, O. A. A. Mohamad, L. Gao, Y.-G. Xie, R. Abdugheni, Y. Huang, L. Li, B.-Z. Fang and W.-J. Li. Sediment prokaryotic microbial community and potential biogeochemical cycle from saline lakes shaped by habitat. Microbiol. Res. 270 (2023), 127342.

[89] Z. Zhao, S. Grohmann, L. Zieger, W. Dai and R. Littke. Evolution of organic matter quantity and quality in a warm, hypersaline, alkaline lake: The example of the Miocene Nördlinger Ries impact crater, Germany. Frontiers in Earth Science 10 (2022).

[90] L. Yue, W. Kong, M. Ji, J. Liu and R. M. Morgan-Kiss. Community response of microbial primary producers to salinity is primarily driven by nutrients in lakes. Sci. Total Environ. 696 (2019), 134001.

[91] X. Kong, Z. Jiang, Y. Zheng, M. Xiao, C. Chen, H. Yuan, F. Chen, S. Wu, J. Zhang and C. Han. Organic geochemical characteristics and organic matter enrichment of mudstones in an Eocene saline lake, Qianjiang Depression, Hubei Province, China. Mar. Pet. Geol. 114 (2020), 104194.

[92] N. Gunde-Cimerman, A. Plemenitaš and A. Oren. Strategies of adaptation of microorganisms of the three domains of life to high salt concentrations. FEMS Microbiol. Rev. 42(3) (2018), 353–375.

[93] A. Oren. Molecular ecology of extremely halophilic Archaea and Bacteria. FEMS Microbiol. Ecol. 39(1) (2002), 1–7.

[94] A. Oren, Ecology of extremely halophilic microorganisms, The biology of halophilic bacteria, CRC Press2020, pp. 25–53.

[95] F. Rodriguez-Valera, Introduction to saline environments, The biology of halophilic bacteria, CRC Press2020, pp. 1-23.

[96] M. Salma and M. K. A. andMehnaz Samina. Osmoadaptation in halophilic bacteria and archaea. Research Journal of Biotechnology Vol 15 (2020), 5.

[97] J. Dröge and A. C. McHardy. Taxonomic binning of metagenome samples generated by next-generation sequencing technologies. Briefings in bioinformatics 13(6) (2012), 646–655.

[98] B. Lobb, B. J.-M. Tremblay, G. Moreno-Hagelsieb and A. C. Doxey. An assessment of genome annotation coverage across the bacterial tree of life. Microbial Genomics 6(3) (2020).

[99] M. Jain, H. E. Olsen, B. Paten and M. Akeson. The Oxford Nanopore MinION: delivery of nanopore sequencing to the genomics community. Genome biology 17(1) (2016), 1–11.

[100] M. Wang, C. R. Beck, A. C. English, Q. Meng, C. Buhay, Y. Han, H. V. Doddapaneni, F. Yu, E. Boerwinkle and J. R. Lupski. PacBio-LITS: a large-insert targeted sequencing method for characterization of human disease-associated chromosomal structural variations. BMC Genomics 16(1) (2015), 1–12.

[101] T. Nawy. Single-cell sequencing. Nat. Methods 11(1) (2014), 18–18.

[102] A. Hoarfrost, A. Aptekmann, G. Farfañuk and Y. Bromberg. Deep learning of a bacterial and archaeal universal language of life enables transfer learning and illuminates microbial dark matter. Nature communications 13(1) (2022), 2606.

[103] J. Gutleben, M. Chaib De Mares, J. D. Van Elsas, H. Smidt, J. Overmann and D. Sipkema. The multi-omics promise in context: from sequence to microbial isolate. Crit. Rev. Microbiol. 44(2) (2018), 212–229.

[104] D. Cui, L. Kong, Y. Wang, Y. Zhu and C. Zhang. In situ identification of environmental microorganisms with Raman spectroscopy. Environmental Science and Ecotechnology (2022), 100187.

[105] W. Zheng, S. Zhao, Y. Yin, H. Zhang, D. M. Needham, E. D. Evans, C. L. Dai, P. J. Lu, E. J. Alm and D. A. Weitz. High-throughput, single-microbe genomics with strain resolution, applied to a human gut microbiome. Science 376(6597) (2022), eabm1483.

[106] K. Yu, S. Yi, B. Li, F. Guo, X. Peng, Z. Wang, Y. Wu, L. Alvarez-Cohen and T. Zhang. An integrated meta-omics approach reveals substrates involved in synergistic interactions in a bisphenol A (BPA)-degrading microbial community. Microbiome 7(1) (2019), 16.

[107] S. Banerjee, A. Bedics, P. Harkai, B. Kriszt, N. Alpula and A. Táncsics. Evaluating the aerobic xylene-degrading potential of the intrinsic microbial community of a legacy BTEX-contaminated aquifer by enrichment culturing coupled with multi-omics analysis: uncovering the role of Hydrogenophaga strains in xylene degradation. Environmental Science and Pollution Research 29(19) (2022), 28431–28445.

[108] W. H. Lewis, G. Tahon, P. Geesink, D. Z. Sousa and T. J. Ettema. Innovations to culturing the uncultured microbial majority. Nature Reviews Microbiology 19(4) (2021), 225–240.

[109] H. Imachi, M. K. Nobu, N. Nakahara, Y. Morono, M. Ogawara, Y. Takaki, Y. Takano, K. Uematsu, T. Ikuta and M. Ito. Isolation of an archaeon at the prokaryote–eukaryote interface. Nature 577(7791) (2020), 519–525.

[110] S. N. Dedysh. Cultivating uncultured bacteria from northern wetlands: knowledge gained and remaining gaps. Frontiers in microbiology 2 (2011), 184.

[111] A. M. Wenger, P. Peluso, W. J. Rowell, P.-C. Chang, R. J. Hall, G. T. Concepcion, J. Ebler, A. Fungtammasan, A. Kolesnikov, N. D. Olson, A. Töpfer, M. Alonge, M. Mahmoud, Y. Qian, C.-S. Chin, A. M. Phillippy, M. C. Schatz, G. Myers, M. A. DePristo, J. Ruan, T. Marschall, F. J. Sedlazeck, J. M. Zook, H. Li, S. Koren, A. Carroll, D. R. Rank and M. W. Hunkapiller. Accurate circular consensus long-read sequencing improves variant detection and assembly of a human genome. Nat. Biotechnol. 37(10) (2019), 1155–1162.

[112] D. M. Bickhart, M. Kolmogorov, E. Tseng, D. M. Portik, A. Korobeynikov, I. Tolstoganov, G. Uritskiy, I. Liachko, S. T. Sullivan, S. B. Shin, A. Zorea, V. P. Andreu, K. Panke-Buisse, M. H. Medema, I. Mizrahi, P. A. Pevzner and T. P. L. Smith. Generating lineage-resolved, complete metagenome-assembled genomes from complex microbial communities. Nat. Biotechnol. (2022).

[113] A. S. Deshpande, N. Ulahannan, M. Pendleton, X. Dai, L. Ly, J. M. Behr, S. Schwenk, W. Liao, M. A. Augello and C. Tyer. Identifying synergistic high-order 3D chromatin conformations from genome-scale nanopore concatemer sequencing. Nat. Biotechnol. 40(10) (2022), 1488–1499.

[114] Y. Huang, R. U. Sheth, S. Zhao, L. A. Cohen, K. Dabaghi, T. Moody, Y. Sun, D. Ricaurte, M. Richardson and F. Velez-Cortes. High-throughput microbial culturomics using automation and machine learning. Nat. Biotechnol. (2023), 1–10.

