## Supplementary figures and images for "Unravelling Biosynthesis and Biodegradation Potentials of Microbial Dark Matters in Hypersaline Lakes"

### Supplemental Figure 1

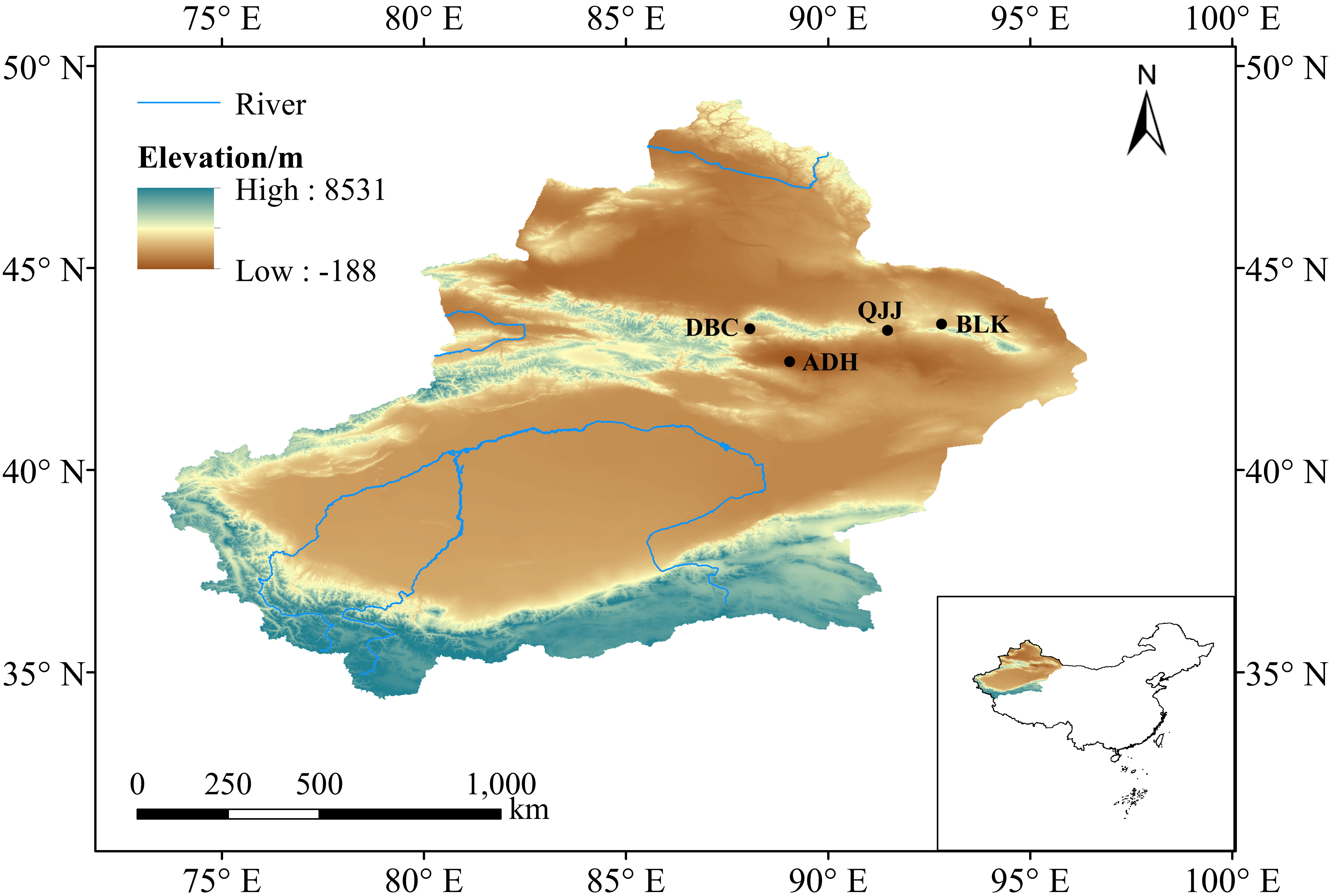

### Supplemental Figure 2

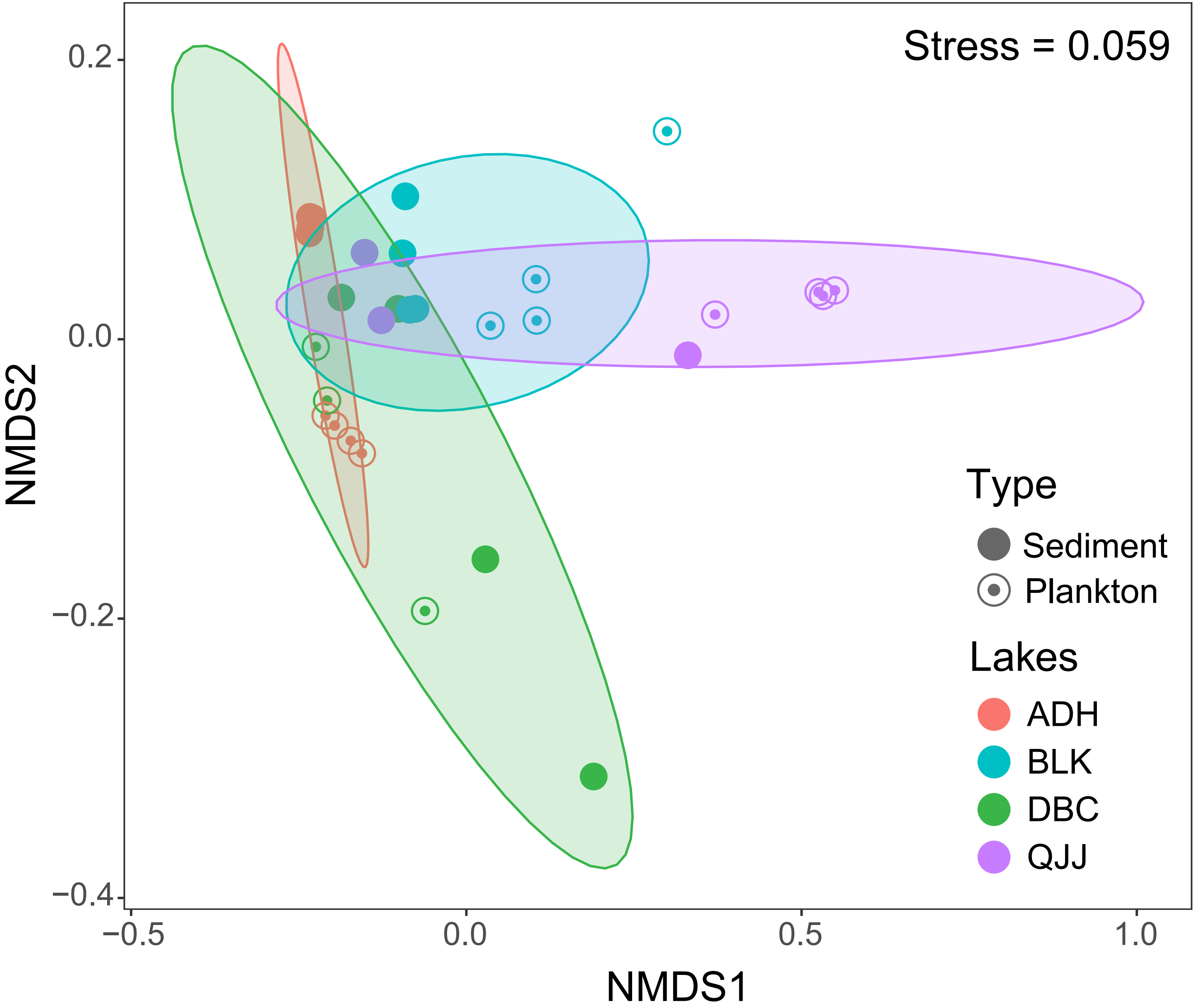
